# Silencing *Doublesex* expression triggers three-level pheromonal feminization in *Nasonia* males

**DOI:** 10.1101/2021.05.21.445141

**Authors:** Yidong Wang, Weizhao Sun, Sonja Fleischmann, Jocelyn G. Millar, Joachim Ruther, Eveline C Verhulst

## Abstract

The transcription factor Doublesex (Dsx) has a conserved function in controlling sexual morphological differences in insects, but our knowledge on its role in regulating sexual behavior is widely limited to *Drosophila*. Here, we show in the parasitoid wasp *Nasonia vitripennis* that males whose Dsx gene had been silenced by RNA interference (*NvDsx*-i) underwent a three-level pheromonal feminization: (1) *NvDsx*-i males were no longer able to attract females from a distance, owing to drastically reduced titers of the abdominal long-range sex pheromone. (2) *NvDsx*-i males were courted by wild-type males like females which correlated with a lower abundance of alkenes in their cuticular hydrocarbon (CHC) profiles. Supplementation of *NvDsx*-i male CHC profiles with realistic amounts of synthetic (*Z*)-9-hentriacontene (*Z*9C31), the most significantly reduced alkene in *NvDsx*-i males, interrupted courtship by wild-type conspecific males. Supplementation of female CHC profiles with *Z*9C31 reduced courtship and mating attempts by wild-type males. These results prove that *Z*9C31 is crucial for sex discrimination in *Nasonia.* (3) *Nvdsx*-i males were hampered in eliciting female receptivity during courtship and thus experienced severely reduced mating success, suggesting that they are unable to produce the hitherto unidentified oral aphrodisiac pheromone reported in *N. vitripennis* males. We conclude that Dsx is a multi-level key regulator of pheromone-mediated sexual communication in *N. vitripennis*. Silencing *Dsx* by RNA interference provides a new avenue for unraveling the molecular mechanisms underlying the pheromone-mediated sexual communication in insects.

## Introduction

Insect sex pheromones play a major role in initiating and guiding the mating process, and in allowing individuals to reliably recognize conspecific mates. In many insect species one sex releases a blend of volatile sex pheromones that transmits a species-specific message to attract conspecifics of the opposite sex (1–3). The attractiveness of the blend is often based on the quantity and ratio of the pheromone components, which may be correlated with nutritional condition, age, or gamete quality of the sender (4, 5). Over shorter distances, sex-specific mixtures of low-volatility, fatty-acid-derived hydrocarbons on the insect cuticle (cuticular hydrocarbons, CHC) play a role in the recognition and mating processes of many insect species (6–10), and species can often be identified by their unique CHC profiles (6, 11, 12). In many insects, the same CHC compounds are shared by both sexes but vary in their relative abundance (13–15), while others have evolved subsets of sex-specific CHC to either attract potential mates or to repel competitors (16, 17).

Because sex pheromone production is sexually dimorphic in nature, it is generally assumed that the sex determination pathway is involved in regulating sex pheromone production. In insects, sexual development is initiated by diverse species-specific primary signals, but downstream, the pathway contains several transcription factors that have relatively conserved functions (18, 19). One major transcription factor, Doublesex (Dsx), is located at the bottom of the cascade, and regulates sexual differentiation in various species across insect orders (20–26). At first, Dsx was considered to be exclusively a regulator of sex-specific morphological development, while the transcription factor Fruitless (Fru), as identified in *Drosophila melanogaster,* was thought to mediate sex-specific behaviours, including mating behaviour. In *Drosophila*, the male Fru protein (Fru^M^) is a key regulator of crucial mating rituals in both the central nervous system (CNS) and the peripheral nervous system (27, 28). Accumulating evidence has been shown that Dsx coordinates the development of the CNS, and together with Fru^M^, shapes courtship behaviour in *D. melanogaster* males (29–31). Nevertheless, Dsx knockdown experiments in other insect species, for example in *Onthophagus taurus* beetles, changed aggressiveness, but did not yield solid evidence for changes in mating behaviours (32). Our knowledge of the role of Dsx in directly regulating the mating behaviour of insects is therefore primarily limited to *Drosophila* flies.

Mating is a complex process in insects, which is influenced by morphological characteristics, behaviour, and chemical communication, making it a challenge to study the role of Dsx in regulating these interacting factors. The jewel wasp *Nasonia vitripennis* (Hymenoptera: Pteromalidae), as an emerging model species, offers great opportunities for this kind of research. *Nasonia vitripennis* is a gregarious parasitoid of several pest fly species (33). The sex determination mechanism in *N. vitripennis* has been completely elucidated (34–37), and recently we illustrated the role of Dsx in regulating sexually dimorphic traits (38). In parallel, the chemical communication system of *N. vitripennis* has been intensively studied (39). The male-derived long-range sex attractant pheromone of *N. vitripennis* consists of the major components (4*R*,5*R*)- and (4*R*,5*S*)-5-hydroxy-4-decanolide (HDL-*RR* and HDL-*RS*) with a synergistic minor compound, 4-methylquinazoline (4MQ) (40, 41). The pheromone is synthesized by males in the rectal vesicle (42) and is deposited onto the natal host and other substrates through a typical “abdominal dipping” behaviour (43, 44). Pheromone marks are highly attractive to conspecific virgin females, whereas mated females are no longer attracted (45). In addition to the male-derived sex attractant, sex- and species-specific CHC profiles have been identified in wasps of the genus *Nasonia* (6, 13, 46, 47). CHC are used for mate recognition, and elicit stereotyped courtship behaviour in *N. vitripennis* males which has been well-characterized and described in detail (48–50). After encountering a female, the male immediately mounts, places his fore tarsi on her head, and starts to perform the mating ritual, which includes stereotypical wing movements, a series of head nods, and antennal sweeping (48). During head nodding, the male moves his mouthparts close to the proximal region of the female’s antennae and synchronizes the extrusion and retraction of his mouthparts with the upward and downward nodding process (48, 49). During this crucial step, males are believed to secrete an aphrodisiac pheromone from the mouthparts onto the female antennae to elicit female receptivity (50). The female signals receptivity by lowering her antennae while raising her abdomen to expose her genital orifice, after which mating usually occurs. Subsequently, another short round of courting, termed post-copulatory courtship, is repeated by the male, during which the female often again exhibits a receptive posture. The sequence terminates with the male dismounting from the female. The blend of oral pheromones apparently not only serves as an aphrodisiac, but also inhibits the response of mated females to the male sex pheromone, thus lowering the chance of re-mating (39). Taken together, many aspects of the *Nasonia* mating behaviour and chemical signaling are known, but the possible role of *Dsx* in regulating these sex- and species-specific traits has not yet been investigated. In this research, we explored the role of *N. vitripennis Dsx* (*NvDsx*) in regulating male mating behaviour and sex pheromone production. We silenced *NvDsx* in male pupae by RNA interference (*NvDsx*-i) and observed each step of the resulting adults’ mating behaviour after emergence. Using coupled gas chromatography-mass spectrometry (GC-MS), we then investigated the effects of *NvDsx* on the pheromone chemistry by comparing long-range sex pheromone titers and CHC profiles of *NvDsx*-i males with those of *green fluorescent protein* RNA interference mock control males (*gfp*-i). Finally, we performed Y-tube olfactometer bioassays and bioassays with treated dummies to determine the attractiveness of *NvDsx*-i males to virgin females with which we could identify key components of the male CHC that inhibit male-male mating attempts.

## Results

### Courting *NvDsx*-i males are less successful in rendering conspecific females receptive

We first asked if the silencing of *NvDsx* expression affects the mating behaviour of males. We microinjected *Dsx* dsRNA into white male pupae and measured a significant suppression of *NvDsx* in these *NvDsx*-i males after emergence when compared to the *gfp*-i control males. (GLM, *P*<0.001), or to Non-injected males (Non-i, GLM, *P*<0.001) (Fig. 1a). In mating trials, we exposed virgin females to *NvDsx*-i, *gfp*-i, or Non-i males and recorded the duration of the behavioural elements searching, mounting including head nodding, copulation, and post-copulatory courtship, as well as male mating success rate. Only 10.9% of *NvDsx*-i males elicited receptivity and subsequently mated, which was significantly less than Non-i (97.6%) and *gfp*-i males (100%, Fig. 1b). *NvDsx*-i males did not differ from Non-i (GLM, *P*=0.940) and *gfp*-i (GLM, *P*=0.951) males with respect to searching times, but the high fraction of unreceptive females led to a significantly longer mounting time for *NvDsx*-i males (Non-i: GLM, *P*<0.001; *gfp*-i: GLM, *P*<0.001, Fig. 1c). The few *NvDsx*-i males that elicit female receptivity after mounting showed no significant difference in copulation time (Non-i: GLM, *P*=0.540; *gfp*-i: GLM, *P*=0.266), but displayed significantly shorter post-copulatory courtship than Non-i males (GLM, *P*<0.001) and *gfp*-i males (GLM, *P*=0.002). There was an effect of *gfp* RNAi on the courtship behaviour, as *gfp*-i males showed longer mounting time (GLM, *P*<0.001) and shorter post-copulatory courtship (GLM, *P*=0.014) than Non-i males. However, the 100% mating success rate of *gfp*-i males indicates overall mating performance was unaffected.

**Fig. 1.**
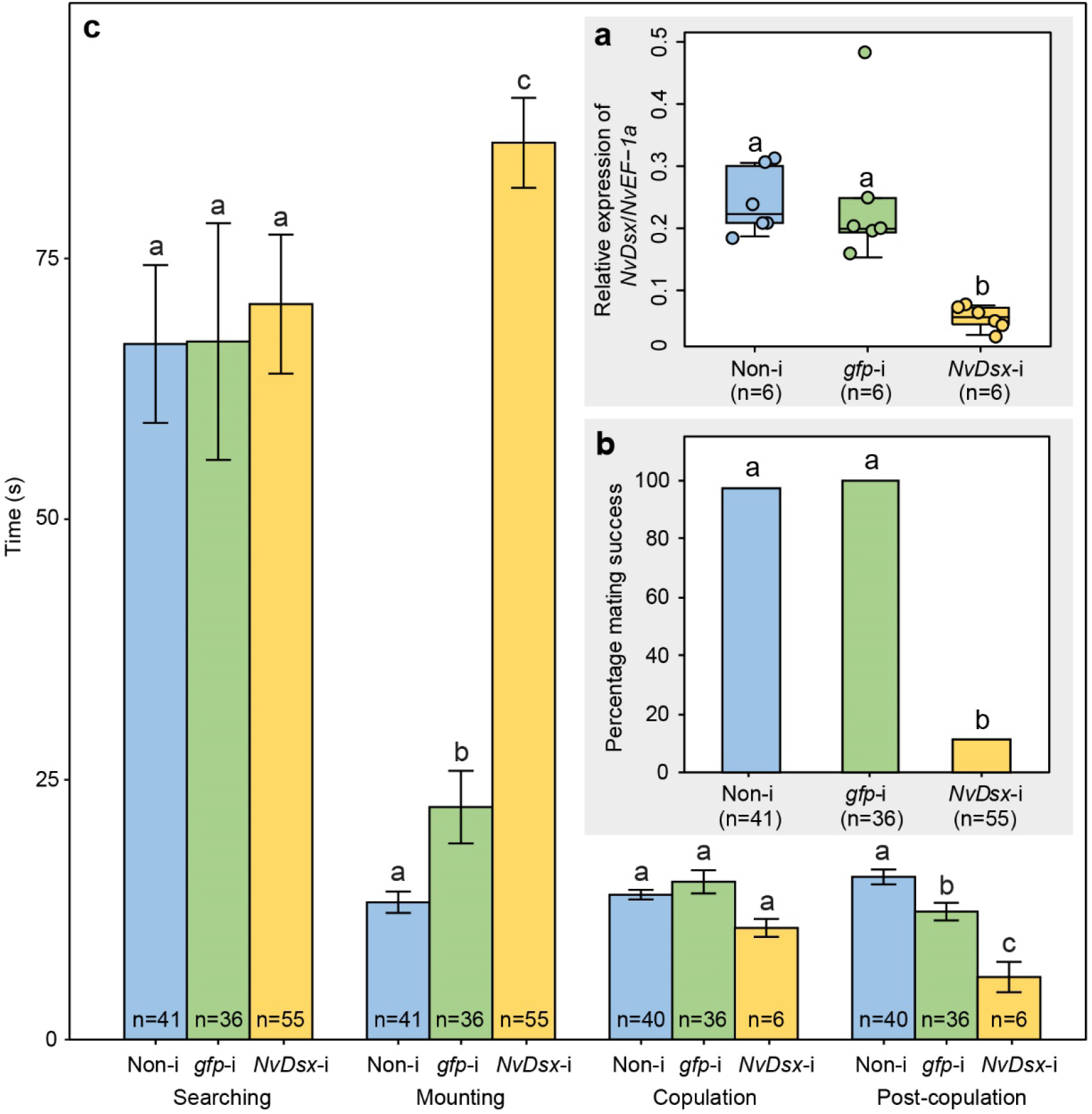
Effects of *NvDsx* knockdown on *NvDsx* expression and mating behaviour of *N. vitripennis* males. **a** Relative expression of *NvDsx/NvEF-1a* in Non-i, *gfp*-i and *NvDsx*-i males. Box-and-whisker plot shows median values (horizontal line), 25-75% quartiles (box), maximum/minimum range (whiskers) and individual data points. **b** Percentage mating success of Non-i, *gfp*-i and *NvDsx*-i males in mating trials with virgin females. **c** Duration of selected behavioural elements (means ± SE) of the *N. vitripennis* mating sequence of Non-i, *gfp*-i, and *NvDsx*-i males in mating trials with virgin females. Different lowercase letters indicate significant differences at *P*<0.05 (**a**, **c**: GLM, **b**: Bonferroni-corrected multiple Chi² tests).

### *NvDsx*-i males are no longer attractive to virgin females due to decreased abdominal sex attractant pheromone production

To evaluate whether *NvDsx*-i males were less successful in attracting virgin females from a distance, we gave virgin females the choice between the odours of two groups of ten live conspecifics in a Y-tube olfactometer. As expected, virgin females preferred the odour of Non-i males (GLM, *P*<0.01, Fig. 2a) or *gfp*-i males (GLM, *P*<0.001) over that of Non-i females. The attractiveness of Non-i and *gfp*-i males did not differ in these trials (GLM, *P*=0.904). In contrast, virgin females were not attracted by the odours of *NvDsx*-i males, showing no preference between those odours and the odours of Non-i females (GLM, *P*=0.856). When given the choice, virgin females did not discriminate between the odours of Non-i males and *gfp*-i males but they preferred Non-i males over *NvDsx*-i males (GLM, *P*<0.001, Fig. 2a). The reduced attractiveness of *NvDsx*-i males in comparison to *gfp*-i males correlated with significantly reduced titres of the abdominal sex attractant pheromones, as determined by GC-MS (Fig. 2b). All three abdominal sex attractant pheromone components, HDL-*RR*, HDL-*RS*, and 4MQ, were detected only in trace amounts in extracts of the abdomens of *NvDsx*-i males, while the total amount and relative proportions of abdominal sex attractant pheromone components in *gfp*-i males were within the range reported previously for Non-i males (41, 51, 52) (Fig. 2c-d).

**Fig. 2.**
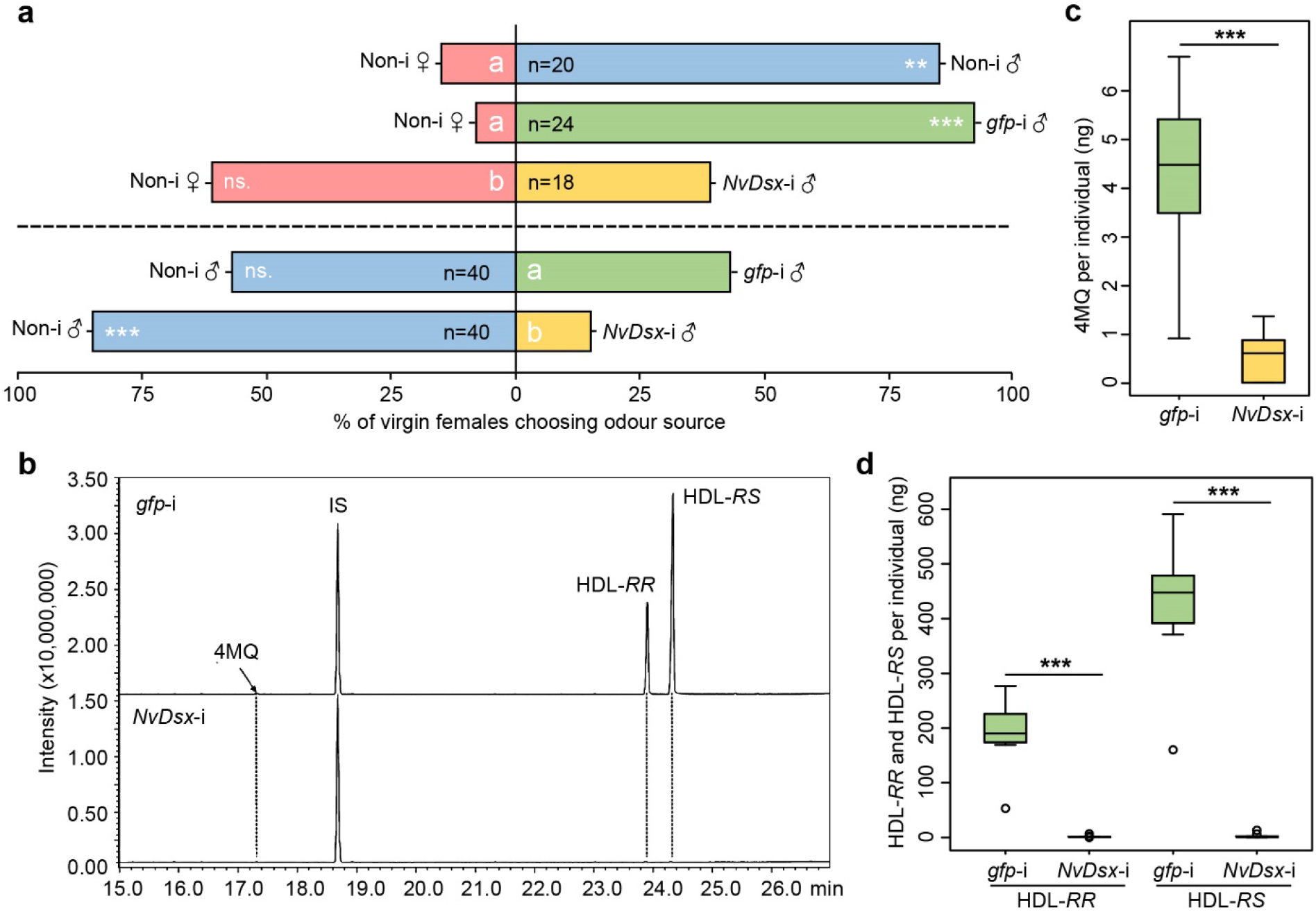
Response of virgin females to the odours of differently treated conspecifics in a Y-tube olfactometer, and chemical analysis of the abdominal sex attractant pheromone in *gfp*-i and *NvDsx*-i males. **a** Percentage of virgin females choosing between the odour of ten live Non-i females, Non-i males, *gfp*-i, or *NvDsx*-i males. The treatments tested in each experiment are given on each side of the bars. Statistical analyses were conducted in two groups separated by the dashed line. Letters indicate significant differences at *P*<0.05 (GLM) and deviations from equality are indicated by asterisks, ns. refers to not significant (*P*>0.05), ** refers to *P*<0.01, *** refers to *P*<0.001 (GLM). **b** Total ion current chromatograms of abdominal extracts from *gfp*-i and *NvDsx*-i males (IS = internal standard, 10 ng methyl undecanoate, HDL-*RR* and HDL-*RS* = (4*R*,5*R*)- and (4*R*,5*S*)-5-hydroxy-4-decanolide, 4MQ = 4-methylquinazoline). Quantification of **c** 4MQ, and **d** HDL-*RR* and HDL-*RS* in abdominal extracts from *gfp*-i and *NvDsx*-i males. Box-and-whisker plots show median (horizontal line), 25-75 % quartiles (box), maximum/minimum range (whiskers) and outliers (° >1.5 × interquartile range outside the first or the third quartile). Asterisks indicate significant differences between treatments at *P*<0.001 (Mann-Whitney *U*-test, n=10).

### *NvDsx*-i males elicit courtship in conspecific males due to decreased alkene abundance in their CHC profiles

Earlier observations of *NvDsx*-i mating behaviour by our group suggested that *NvDsx*-i males are mistaken for females by normal males. Previous research also showed that males use CHC signaling to discriminate between the sexes, and that female-derived CHC arrests males and elicits courtship behaviour (13). Hence, we hypothesized that the unusual courting of *NvDsx*-i males by Non-i males is elicited by changes in their CHC profiles. One characteristic feature of *N. vitripennis* CHC profiles is the male-biased relative abundance of alkenes (13). Analysis of the CHC profiles of *NvDsx*-i and *gfp*-i males indeed revealed a significant reduction in alkenes (Fig. 3a-b). The amounts of (*Z*)-9-hentriacontene (*Z*9C31) in CHC extracts from *NvDsx*-i males were less than one-fourth of those found in *gfp*-i males (mean ± SE *gfp*-i: 31.8 ± 5.7 ng; *NvDsx*-i: 6.6 ± 1.1 ng). Similar reductions, albeit to a lesser extent, were found for (*Z*)-7-hentriacontene (*Z*7C31), (*Z*)-9-tritriacontene (*Z*9C33), and (*Z*)-7-tritriacontene (*Z*7C33).

**Fig. 3.**
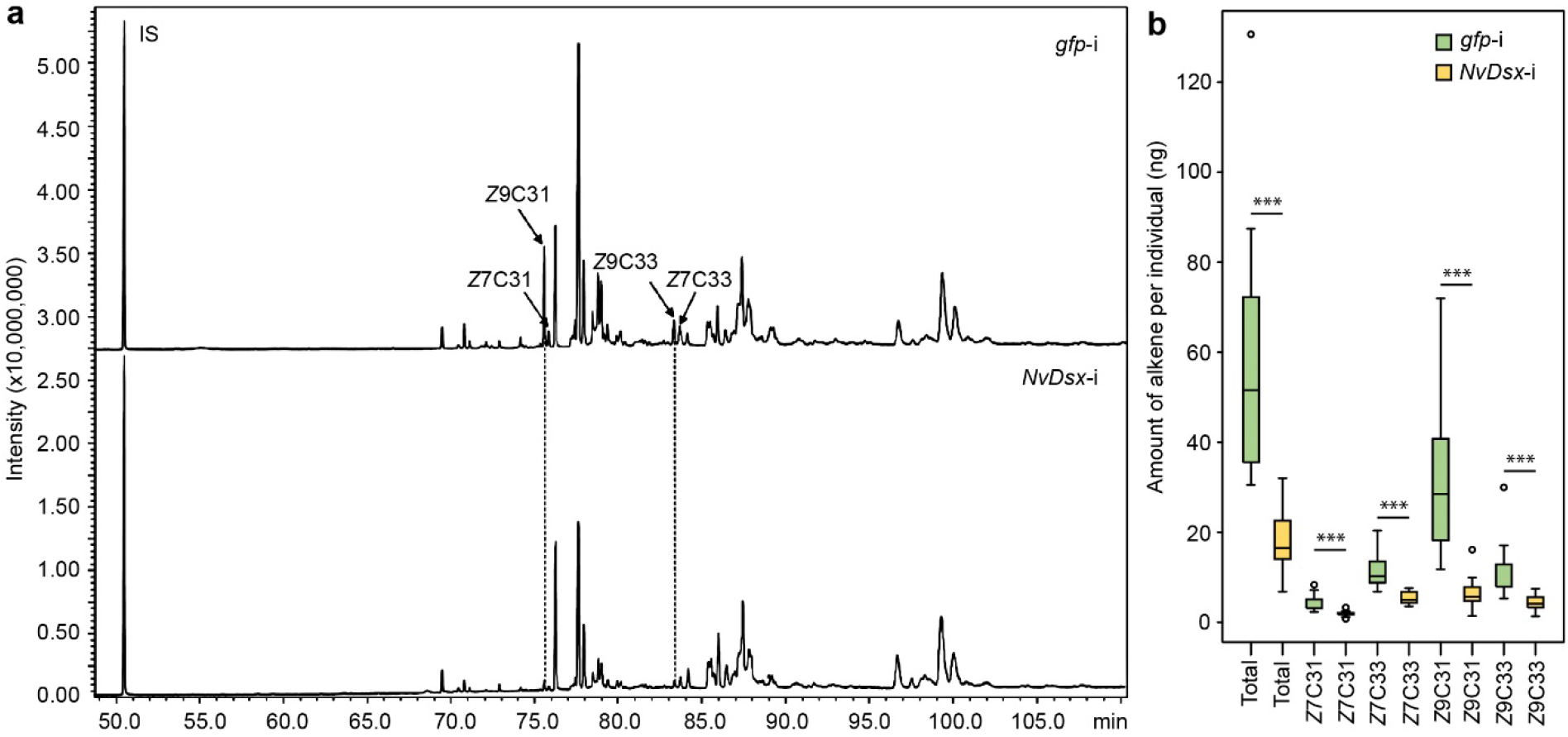
Chemical analyses of cuticular hydrocarbons in *NvDsx*-i and *gfp*-i males. **a** Total ion chromatograms of whole-body hexane extracts from *gfp*-i and *Nvdsx*-i males. Arrows indicate peaks of the alkenes (*Z*)-9- and (*Z)*-7-hentriacontene (*Z*9C31 and *Z*7C31) as well as (*Z*)-9- and (*Z*)-7-tritriacontene (*Z*9C33 and *Z*7C33). IS = internal standard (10 ng tetracosane). **b** Quantification of alkenes in dichloromethane extracts of abdomens of *gfp*-i and *Nvdsx*-i males. The box-and-whisker plots show median (horizontal line), 25-75 % quartiles (box), maximum/minimum range (whiskers) and outliers (° >1.5 × interquartile range outside the first or third quartile). Asterisks indicate significant differences between treatments at *P*<0.001 (Mann-Whitney *U*-test, n = 10).

These results suggest that the decrease in alkenes explains the same-sex courtship of *NvDsx*-i males. Given that *NvDsx*-i treatment affected the amounts of *Z*9C31 the most, we tested whether the application of natural amounts of this compound to *NvDsx*-i males was sufficient to interrupt this male-male courtship. *Nasonia* males are known to respond to attractive dead conspecifics by mounting and copulation attempts (50). Therefore, we conducted a series of behavioural experiments with differently treated dead wasps (dummies). We presented Non-i males with untreated *gfp*-i and *NvDsx*-i dummies as well as with those treated with 30 ng of synthetic *Z*9C31 dissolved in dichloromethane (DCM) or the pure DCM solvent (control). In general, a higher percentage of *N. vitripennis* Non-i males tried to copulate with *NvDsx*-i male dummies than with *gfp*-i male dummies (*X^2^*(1, *N*=50)=8, *P*=0.028, Fig. 4a), spent significantly more time on *NvDsx*-i male dummies than on *gfp*-i male dummies (Mann-Whitney *U*=149, *n*_1_=*n*_2_=25, *P*<0.01, Fig. 4b), and tried to copulate with *NvDsx*-i male dummies longer than with *gfp*-i male dummies (Mann-Whitney *U*=209, *n*_1_=*n*_2_=25, *P*=0.012, Fig. 4c). The application of pure DCM to *NvDsx*-i male dummies did not significantly alter the courtship behaviour of Non-i males (Fig. 4a-c). However, comparing to the pure DCM solvent treated *NvDsx*-i male dummies, the application of *Z*9C31 to *NvDsx*-i male dummies significantly reduced the proportion of Non-i males attempting to copulate with those dummies (*X^2^*(1, *N*=50)=20.053, *P*<0.001), the duration of mounting (Mann-Whitney *U*=46.5, *n*_1_=*n*_2_=25, *P*<0.001) and copulation attempts (Mann-Whitney *U*=130, *n*_1_=*n*_2_=25, *P*<0.01), to the levels of *gfp*-i dummies (Fig. 4a-c). These results support our hypothesis that males use the increased abundance of alkenes in the CHC profiles of males to discriminate between sexes. Therefore, in the next step we investigated whether treatment with *Z*9C31 would make female dummies less attractive to responding males. To this end, we performed a similar dummy experiment as described above, using Non-i male (negative control) and differently treated Non-i female dummies. A significantly higher proportion of *N. vitripennis* Non-i males tried to copulate with Non-i female dummies than with Non-i male dummies (*X^2^*(1, *N*=40)=28.972, *P*<0.001), and Non-i males mounted (Mann-Whitney *U*=22, *n*_1_=*n*_2_=20, *P*<0.001) and tried to copulate (Mann-Whitney *U*=21, *n*_1_=*n*_2_=20, *P*<0.001) significantly longer with Non-i female dummies than with Non-i male dummies (Fig. 4d-f). Application of pure DCM solvent to Non-i female dummies did not influence the response of Non-i males. However, comparing to the pure DCM solvent treated Non-i female dummies, application of 30 ng of *Z*9C31 to Non-i female dummies reduced the proportion of mating attempts by Non-i males (*X^2^*(1, *N*=40)=13.333, *P*<0.01) as well as the total time Non-i males spent trying to copulate with the Non-i female dummies to intermediate levels (Mann-Whitney *U*=59, *n*_1_=*n*_2_=20, *P*<0.001, Fig. 4d,f). The duration of mounting was also reduced and was not significantly different than that of Non-i male dummies (Mann-Whitney *U*=121, *n*_1_=*n*_2_=20, *P*=0.202, Fig. 4e).

**Fig. 4.**
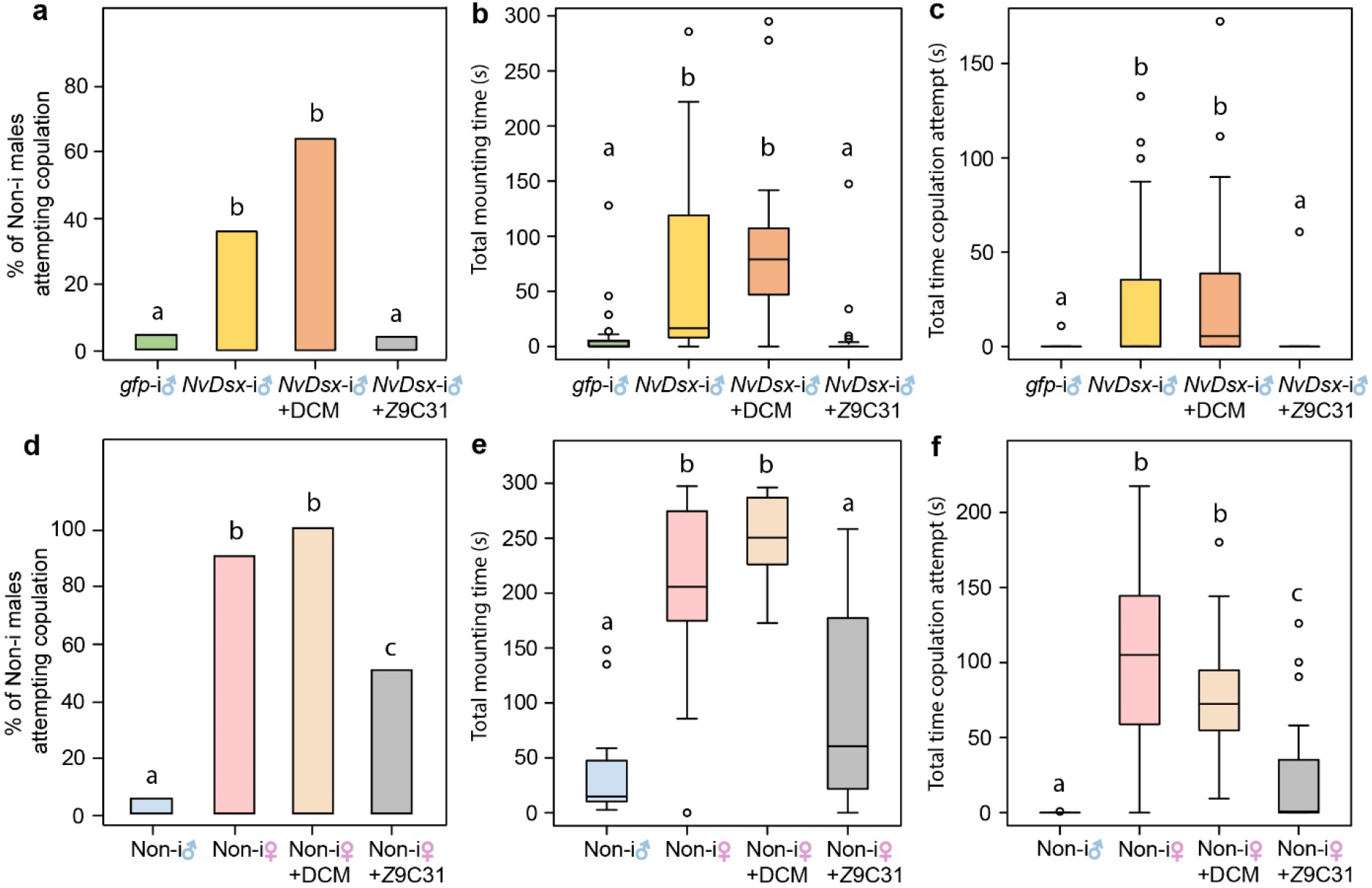
Effect of synthetic *Z* 9C31 on the courting behaviour of Non-i *N. vitripennis* males. Responding Non-i males were presented with dead conspecifics (dummies) for a five minutes observation period. Dummies were either differently treated *gfp*-i or *NvDsx*-i males (a-c, n=25) or Non-i wasps of either sex (d-f, n=20). Tested dummies were either untreated, treated with 30 ng of (*Z*)-9-hentriacontene (*Z*9C31) dissolved in dichloromethane (DCM), or with pure DCM. Shown is **a**, **d** the percentage of Non-i males trying to copulate with the dummy, **b**, **e** the total mounting time, and **c**, **f** the duration of copulation attempts. Box-and-whisker plots (b, c, e, and f) show median (horizontal line), 25-75 % quartiles (box), maximum/minimum range (whiskers) and outliers (° >1.5 × interquartile range outside the first or third quartile). Letters indicate significant differences at *P*<0.05 (data analysis by Kruskal-Wallis H-test followed by Bonferroni-corrected multiple Mann-Whitney *U*-tests (**b**, **c**, **e**, and **f**) or by Bonferroni-corrected multiple 2×2 Chi^2^-tests (**a** and **d**).

## Discussion

In this study we set out to identify the role of Dsx in regulating *N. vitripennis* male mating behaviour and pheromone production, and we found a major effect of NvDsx in the regulation of the production of both volatile and non-volatile sex pheromones. Using Y-olfactometer two-choice assays and GC-MS analyses, we observed a reduced attractiveness of *NvDsx*-i males to virgin females accompanied by the almost complete disappearance of the abdominal volatile sex attractant pheromone components HDL-*RR*, HDL-*RS*, and 4MQ. HDL-*RR* and HDL-*RS* are synthesized by *N. vitripennis* males in the rectal vesicle from fatty acids, and are released via the anal orifice (42). Our results suggest that NvDsx directly regulates the biosynthetic pathway, but which of the genes involved in the biosynthesis of HDL (53) are targeted by NvDsx, remains unknown. Although it has been suggested that the production of the CHC pheromones of females is controlled by Dsx in *D. melanogaster* (54, 55), hardly any research has focused on this topic in other insects. Hence, the present study is, to our knowledge, the first to demonstrate a regulatory effect of Dsx on the biosynthesis of pheromones other than CHC in insects.

Despite their lack of abdominal sex attractant pheromones for long-range attraction of females, all tested *NvDsx*-i males were still able to mount and court females, and displayed stereotypical mating behaviours such as head nodding and antennal sweeping. Apparently, the substantial reduction in the titres of the sex attractant pheromone did not affect the acceptance of courtship by females, initiated by the males through mounting. We also found no significant differences in the searching times for the different treatments in the mating experiment, which may have been due to the experimental set-up, in which the male and female were enclosed in a small space. At this level of the mate finding process, the abdominal sex attractant pheromone of *N. vitripennis* males no longer plays a role, and it is up to the male to recognize and court the female via the female-specific CHC (13). Intriguingly, *NvDsx*-i males were courted by Non-i males in a similar manner to females, suggesting a kind of “feminization” of the *NvDsx*-i CHC profiles. *NvDsx*-i males apparently became sexually attractive to Non-i males because of a substantial reduction of alkenes, in particular *Z*9C31, in their CHC profile. Supplementation of the profile by applying synthetic *Z*9C31 to *NvDsx*-i male dummies eliminated the courting behaviour of Non-i males, suggesting that *Z*9C31 is a key compound for sex discrimination by *N. vitripennis* males. Just as the cuticular pheromone (*Z*)-7-tricosene inhibits male-male courtship in *D. melanogaster* (16, 56)*, Z*9C31 helps to prevent male-male courtship in *N. vitripennis*. The low abundance of *Z*9C31 in the CHC profile of females (13) might even underlie males’ recognition of potential mates. This idea is supported by our finding that applying *Z*9C31 to female dummies reduced Non-i male courting attempts and duration, although not to the low levels shown towards Non-i males. In this context, it is important to note that the other three alkenes, and many other CHC components that are found in different relative amounts in *N. vitripennis* males and females might also contribute to this effect (13, 57). Hence, the addition of *Z*9C31 alone might not be sufficient to entirely disrupt a female’s pheromone signal.

We analyzed the presence and amounts of alkenes that we found to be directly or indirectly regulated by NvDsx in the previously published, species-specific CHC profiles of *Nasonia* (6). We found a male bias in the expression of these alkenes in *N. vitripennis* but not in the other three *Nasonia* species (6, 47). In *D. melanogaster,* alkenes are a major fraction of the CHC profiles (58), and their biosynthesis requires different types of desaturases (59). Niehuis et al. (2011) found that two of the *Drosophila* desaturase gene homologs map to regions in the *Nasonia* genome that contain the QTL clusters for these alkenes. NvDsx is a prime candidate for regulation of the expression of the desaturase genes that controls alkene biosynthesis. The gain and loss of the binding-site of Dsx in these desaturase genes may account for the species-specific differences in alkene synthesis, as has been shown for *D. melanogaster* (60). This difference might be related to specific life history traits of different *Nasonia* species. Females mating inside the host puparium (within-host mating, WHM) is rare in parasitoids, but it does happen in *Nasonia* (61). Compared to *N. vitripennis,* the sibling species *N. giraulti* and *N. longicornis* show a significantly higher WHM rate (61). Because *N. vitripennis* is ancestral to the other species in the genus (62), this sex-specific difference in alkene expression might have been lost in *N. giraulti* and *N. longicornis* as a result of decreased mate competition due to their higher WHM rates. Further research will be required to verify this. In addition, behavioural assays to determine mate discrimination among *Muscidifurax uniraptor, N. vitripennis*, and *Trichomalopsis sarcophagae* showed that *N. vitripennis* males do not discriminate against *T. sarcophagae* females, displaying courtship behaviour and copulation attempts, but reject *M. uniraptor* females (46). Despite the general similarity of the CHC profiles of females of both species, both *Z*9C31 and *Z*9C33 are produced by *N. vitripennis* and *T. sarcophagae* females at low levels, whereas *M. uniraptor* females have higher amounts of these alkenes in their CHC profiles (46). Hence, *Z*9C31 and the other alkenes might also be involved in species discrimination by *N. vitripennis* males. It would be interesting to further investigate the consequences and benefits of maintaining this alkene-dependent olfactory recognition system.

By comparing the courtship and copulation behaviours of *NvDsx*-i, *gfp*-i, and Non-i males when presented with live virgin females, we did not find obvious differences in male courtship performance between *NvDsx*-i and Non-i males. Male courtship behaviour in insects is thought to be governed primarily by Fruitless (Fru) (28). However, in *Drosophila*, Dsx was shown to facilitate the formation of the CNS, and these neuronal networks develop during pupation (63). This suggests that silencing *NvDsx* from the pupal stage onwards does not affect the development of the regions of the CNS that control mating behaviour in *N. vitripennis* males. Hence, our results add to the current theory that Fru is the main regulator of courtship behaviour in insects, but it should be noted that silencing *NvDsx* earlier in development might produce different outcomes.

We observed significantly prolonged mounting time and a strong reduction in mating success of *NvDsx*-i males when they courted normal females, indicating that an important component inducing receptivity of females might be missing from the *NvDsx*-i males. It has been shown that males with sealed mouthparts cannot induce receptivity in females, which indicates the existence of an aphrodisiac produced by the mouthparts (50, 64). The identity of this aphrodisiac is still unknown despite extensive efforts to unravel its chemical structure (39). So far, three fatty acid ethyl esters have been identified in the oral secretion of *N. vitripennis* males, but these compounds were unable to elicit female receptivity (65). The inability of *NvDsx*-i males to elicit receptivity in conspecific females suggests that the production of the oral aphrodisiac is affected by Dsx, possibly in combination with, or due to malformed mandibular glands, which are known to be present in *N. vitripennis* (66). Therefore, comparative chemical analyses of the heads of *NvDsx*-i and Non-i males will be a promising approach to unravel the identity of the oral aphrodisiac, as a next step towards a comprehensive understanding of pheromone communication in *N. vitripennis*.

## Material and methods

### 1. Insect rearing

The *Wolbachia*-free AsymCx laboratory strain of *N. vitripennis* was used in all experiments. The laboratory cultures were constantly reared on the host *Calliphora sp.* and *Lucilia caesar* (only the wasp for dead wasp bioassay) at 25°C with a 16h/8h light/dark cycle.

### 2. *NvDsx* dsRNA preparation

*NvDsx* knockdown was conducted by using RNA interference (RNAi) as described in Wang et al. (2020). MEGAscript RNAi Kit (Invitrogen™, Waltham, Massachusetts, USA) was used to produce the dsRNA that targets the common region (exons 2-5) of *NvDsx* in all male splice-variants. *Gfp* dsRNA was used as a control for all experiments and was generated from the vector pOPINEneo-3C-GFP (Addgene plasmid # 53534; http://n2t.net/addgene: 53534; RRID: Addgene_53534). Primers that were used to construct the dsRNA T7 template were designed in Geneious 10.0.9 (Table S1). Both *NvDsx* and *gfp* dsRNAs were diluted in RNase-free water to a final concentration of 4000 ng/ul (determined using a NanoDrop™ 2000 Spectrophotometer, Thermo Scientific™) and stored at −20°C.

### 3. Microinjection

DsRNAs were pre-mixed with a prepared red food dye at a 9:1 ratio before microinjection. Commercially available red food dye was prepared by diluting it 2X in DNase/RNase free water. Because asexual reproduction leads to all-male broods in the haplodiploid sex determination system of Hymenoptera, virgin females were used to produce the male offspring that were used at the pupal stage for microinjection. Thirty female pupae were collected in the black pupal stage (~12 days after egg laying), transferred to a single glass vial (7.5 cm length, one cm diameter), plugged with a cotton plug, and kept under rearing conditions as described above. Twenty-four hours after eclosion, each female was provided with two *Calliphora sp.* host pupae per day. Host pupae were continuously incubated under rearing conditions until parasitoid offspring reached the white pupal stage (~7 days after egg laying). Pupae of male *N. vitripennis* were subsequently collected from hosts and fixed on glass slides with double-sided adhesive tape (3M). Microinjection was performed with a FemtoJet^®^ 4i microinjector (Eppendorf, Hamburg, Germany) following the protocol described by Lynch and Desplan (2006). Phosphate Buffered Saline (PBS) plates were prepared by adding 1.5 g BD Bacto™ Agar (Fisher Scientific, Sparks, Nevada, USA) and one Oxoid™ phosphate buffered saline tablet (Thermo Scientific™, USA) into 100 ml distilled water and autoclaving at 115°C for 15 minutes before pouring them into plastic petri-dishes. After microinjection, slides with injected pupae were placed on PBS plates to prevent dehydration and incubated under rearing conditions until adult emergence.

### 4. Quantification of *NvDsx* expression by qPCR

*NvDsx*-i males were collected once they emerged. Six individuals were pooled as one biological replicate and six biological replicates per treatment were collected in 1.5 ml microcentrifuge tubes and flash frozen in liquid nitrogen. ZR Tissue & Insect RNA MicroPrep™ (Zymo Research Corp., Irvine, California, USA) kit was used under the manufacturer’s instructions to extract the total RNA with an on-column DNase treatment step. Sixteen μl of DNase/RNase free water was applied to elute each sample. RNA concentration was determined with a NanoDrop™ 2000 Spectrophotometer (Thermo Scientific™) and one μg of total RNA from each sample was converted into cDNA with SensiFAST™ cDNA Synthesis Kit (Bioline, London, UK) using a T100™ Thermal Cycler (Bio-Rad) with an incubation program consisting of five minutes priming at 25°C, 30 minutes at 46°C and five minutes at 85°C.

Following the manufacturer’s instruction, a SensiFAST™ SYBR^®^ No-ROX Kit (Bioline, London, UK) was used to perform the qPCR. cDNA templates were further diluted 1:100 and five μl aliquots of each diluted template were used in qPCR reactions to verify gene expression. *Elongation factor 1α* of *N. vitripennis* (*NvEF-1α*) was used as a reference gene. Primers that were used to amplify the common region of *NvDsx* in all male splice-variants were designed in Geneious 10.0.9. Detailed primer sequences are listed in Table S1.

### 5. Mating trials

In the mating trials, non-injected (Non-i) females were presented with Non-i males (n=40), *gfp*-i males (n=40), or *NvDsx*-i males (n=60). One- to two-day-old virgin males and females for the mating trials were separated at black pupal stage by sex-specific traits such as forewing size and presence/absence of the ovipositor, and were collected in separate glass vials (7.5 cm length, one cm diameter) plugged with cotton until eclosion. Each wasp was used only once. A virgin female was transferred into a glass vial (7.5 cm length, one cm diameter) that contained a male, and the wasps were brought in contact at the bottom by quickly flicking the vial. Observations started immediately after transferring the female. The durations of four distinctive elements of the mating process were recorded: (1) searching time, which is the time from the start of the observation until the male mounts the female; (2) mounting time, the duration of the male mounting the female; (3) copulation time, the duration of the copulation; (4) post-copulation time, the duration of the male remounting the female after copulation until the male dismounts. Recordings were excluded if mounting did not occur within five minutes. Mating success was pronounced positive if all four mating steps were finalized in a single mating trial and was used to calculate the mating success rate.

### 6. Y-tube olfactometer bioassay

The responses of virgin females to the odours of differently treated conspecifics were tested in a glass still-air Y-tube olfactometer (0.9 cm diameter), consisting of an 8.5 cm stem and two five cm arms. Prior to the tests, ten wasps of either treatment were kept in five ml transparent pipette tips that were sealed at both ends with parafilm. Wasp odours were allowed to accumulate in the pipette tips by keeping the wasps inside the pipette tips for two days under rearing conditions. Afterwards, the bottom parafilm was replaced by a piece of fibre mesh, and pipette tips were mounted to the arms of the Y-tube olfactometer. Prior to each experiment, odours were allowed to diffuse for five minutes from the pipette tips into the arms of the Y-tube olfactometer. Tests started once a glass vial containing a one- to two-day-old virgin female was connected to the stem of Y-tube olfactometer. Tested females were given three minutes to enter the stem of the Y-tube olfactometer and another three minutes to make a decision. Replicates were excluded when females did not enter the stem of the Y-olfactometer in three minutes. A decision was recorded, if a female entered one arm of the olfactometer and stayed less than one cm away from the respective mesh for one minute. Any other cases were recorded as “no choice”. After each test, the Y-olfactometer was turned by 180° to compensate for any unforeseen asymmetry bias. The Y-tube olfactometer was replaced by a clean one after five tests and pipette tips with wasps were replaced by new ones after 10 tests. Each responding female was tested only once and tested female was gently brushed out of the Y-tube. Tested combinations of two choices were between: 10 Non-i females and 10 Non-i males (n=20), 10 Non-i females and 10 *gfp*-i males (n=25), 10 Non-i females and 10 *NvDsx*-i males (n=25), 10 Non-i males and 10 *gfp*-i males (n=40), and 10 Non-i males and 10 *NvDsx*-i males (n=40).

### 7. Behavioural experiments with dead wasps

Experiments were performed in a round bioassay chamber (10 mm diameter × 3 mm high) made from acrylic glass and covered by a cover slip (68). Behaviours were observed by using a stereo microscope with illumination from a microscope light, and recorded using The Observer XT software (Noldus Information Technology, Wageningen, The Netherlands). Non-i males were exposed to differently treated, freeze-killed one- to two-day-old cadavers (dummies) for five minutes. For each test, the time was recorded that the males spent mounting the dummy, whether these males tried to copulate with the dummy and, if so, the time that these copulation attempts lasted. In the first experiment, we tested the responses of Non-i males (n=25) to untreated male *gfp*-i and *NvDsx*-i dummies as well as to male *NvDsx*-i dummies treated with one μl dichloromethane (DCM, control) or one μl of a solution of *Z*9C31 in DCM (30 ng/μl). In the second experiment, we tested the responses of Non-i males (n=20) to untreated Non-i male and female dummies as well as to Non-i female dummies treated with one μl DCM (control) or one μl of a solution of *Z*9C31 in DCM (30 ng/μl). *Z*9C31 was synthesized based on Saul-Gershenz and Millar (2006). The dose of *Z*9C31 was based on the amount that was found in the CHC extracts of *gfp*-i males (see results). In all treatments that used DCM, the solvent was allowed to evaporate for one minute before the start of the bioassays.

### 8. Chemical analyses

For CHC analyses, individual freeze-killed two-day-old *gfp*-i and *NvDsx*-i males (n = 10) were extracted for 30 minutes with 15 μl hexane containing 10 ng/μl tetracosane as an internal standard. For analysis of the volatile sex attractant pheromone (HDL), dissected abdomens of individual freeze-killed two-day-old *gfp*-i and *NvDsx*-i males (n = 10) were extracted with 30 μl of DCM containing 10 ng/μl methyl undecanoate as an internal standard. Aliquots of two μl of extracts (HDL analyses: one μl) were analysed on a Shimadzu QP2010 Plus GC/MS system equipped with a non-polar BPX5 capillary column (60 m × 0.25 mm inner diameter, 0.25 μm film thickness; SGE Analytical Science Europe, Milton Keynes, UK). Samples were injected at 300°C in splitless mode using an AOC 20i auto sampler. Helium was used as carrier gas at a linear velocity of 30 cm s^−1^ (HDL: 37.8 cm s^−1^). The initial oven temperature of 150°C (HDL: 80°C) was increased at 2°C min^−1^ (HDL: 5°C min^−1^) to 300°C (HDL: 280°C) and held for 30 minutes. The mass spectrometer was operated in the electron ionization mode at 70 eV and the mass range was *m/z* 35-600. Identifications of compounds were done by comparing mass spectra and linear retention indices with those of authentic reference chemicals (HDL-*RR*, HDL-*RS*, 4MQ, *Z*9C31) or previously published identifications of specific CHC in this species (13) (*Z*7C31, *Z*9C33, *Z*7C33). Quantification of alkenes and HDL was done by comparing the peak areas of the analytes to those of the respective internal standards.

### 9. Data analysis

qPCR data was first imported to LinRegPCR software (LinRegPCR, 2017.1.0.0, HFRC, Amsterdam, The Netherlands). After baseline correction, the initial number of templates (N0) was calculated based on averaging PCR efficiency in each amplicon. Relative expression levels of target genes were obtained by dividing the N0 value of target genes by the N0 value of the reference gene. Statistical analysis of qPCR and mating behaviour data was performed in R (70) using general linear models (GLM) with gamma distribution and Tukey’s Honestly Significant Difference (HSD) test for post-hoc comparisons. Mating success rates were compared by Bonferroni-corrected multiple 2×2 Chi^2^ tests. Y-tube olfactometer data were analysed in R using GLM with binominal distribution with cloglog link function and HSD for post-hoc comparisons. The amount of alkenes in the CHC extracts and HDL and 4MQ in the abdomen extracts of *gfp*-i and *NvDsx*-i males were compared by Mann-Whitney *U*-tests. In tests with dummies, the total mounting times and the durations of copulation attempts were compared by a Kruskal-Wallis H-test followed by Bonferroni-corrected multiple Mann-Whitney *U*-tests for pairwise comparisons. The percentages of males showing copulation attempts were compared by Bonferroni-corrected multiple 2×2 Chi^2^ tests.

## Data availability

The datasets generated during and/or analysed during the current study are available in the figshare repository, [10.6084/m9.figshare.14572671].

## Author contributions

Y.W. generated the dsRNA. Y.W and W.S. performed microinjections. W.S. conducted mating behaviour bioassays and Y-tube olfactometer experiments. Chemical analyses were carried out by J.R. Bioassays with dummies were performed by S.F. J.M. synthesized the *Z*9C31. Y.W. and J.R. contributed to the data analysis. Y.W., E.C.V. and J.R. contributed to the design of the work. Y.W. drafted the manuscript. E.C.V., J.R., and J.M. revised the draft manuscript.

## Competing interests

The authors declare no competing interests.

## Acknowledgment

We thank Ray Owens for providing the vector pOPINEneo-3C-GFP to support this research.

## Notes

https://figshare.com/articles/dataset/raw_Data_for_NC_xlsx/14572671

### Competing Interest Statement

The authors have declared no competing interest.

