## Supplemental Table S1 for "Silencing *Doublesex* expression triggers three-level pheromonal feminization in *Nasonia* males"

**Supplementary**

**Table S1. Primers used in the paper**

a. Used in Verhulst et al. (2010)

* T7 promoter sequence that is recommended by the manufacturer.

| Primer | Sequence |
| --- | --- |
| Nv_Dsx_qPCR_F^a^ | CAGCAACACGAGATGCTGATGG |
| Nv_Dsx_qPCR_R^a^ | TGTCATTACTGCCATTCATGCTTGG |
| Nv_EF-1a_qPCR_F^a^ | CACTTGATCTACAAATGCGG |
| Nv_EF-1a_qPCR_R^a^ | GAAGTCTCGAATTTCCACAG |
| Nv_Dsx_RNAi_F | *[TAATACGACTCACTATAGGG]CCAAGAGGCAGCAAATTATG |
| Nv_Dsx_RNAi_R | *[TAATACGACTCACTATAGGG]GTTATACGCCGCATGGCTAC |
| GFP_RNAi_F | *[TAATACGACTCACTATAGGG]GTGACCACCTTGACCTACG |
| GFP_RNAi_R | *[TAATACGACTCACTATAGGG]TCTCGTTGGGGTCTTTGCT |
